# The Role of the Posterior Medial Network in Language Comprehension: Dissociating Construction of Episodic versus Semantic Representations

**DOI:** 10.1101/2022.09.03.506471

**Authors:** Francesca M. Branzi, Matthew A. Lambon Ralph

## Abstract

Language comprehension involves the construction of complex mental representations, i.e., “event representations”, reflecting current events or situation models. The construction of these representations requires manipulation of both semantic and episodic content and has been widely associated with the functioning of the posterior medial network, a subsystem of the default network. However, it is still unknown the extent to which activity in posterior medial network reflects construction of the variable episodic *versus* semantic content of event representations. In this fMRI study, we establish the unique neural correlates of (1) episodic richness and (2) semantic coherence measured during a narrative reading task. Interestingly, we observed a functional fractionation within the posterior medial network in terms of brain regions whose activity was modulated by semantic or episodic content. Specifically, the number of episodic details in the narratives modulated the activity in the left angular gyrus and the retrosplenial cortex/ventral posterior cingulate cortex. Semantic coherence, in contrast, modulated neural responses in the right anterior temporal lobe/middle temporal gyrus, but also in brain regions within the posterior medial network, such as the dorsal posterior cingulate cortex. These results provide the first demonstration of functional dissociations within the posterior medial network in terms of brain regions involved in the construction of semantic *versus* episodic representational content during a language comprehension task.

**Significance Statement:** The construction of “event representations”, which is crucial to understand the world around us, predict the future and make plans, requires manipulation of both semantic and episodic information. The functioning of the posterior medial network has been tightly linked to formation of event representations. However, it is unclear the extent to which activity in this network reflects construction of the variable episodic *versus* semantic content of event representations. The present study provides the first demonstration of functional dissociations within posterior medial network in terms of brain regions involved in construction of semantic *versus* episodic representational content during language comprehension. These findings represent a first step towards understanding how episodic and semantic memory systems operate during the construction of event representations.

## 1. Introduction

The construction of “event representations” requires manipulation of both episodic and semantic knowledge (Bailey et al., 2017). During language comprehension, incoming information is continuously compared and integrated with the current internal “situational model” that (1) specifies how different entities in the environment are related one to another (Zwaan and Radvansky, 1998) and (2) is informed by semantic knowledge (Zwaan et al., 1995) assimilated from previous experiences about elements contained within the situation as well as their associations (Binder and Desai, 2011).

The formation of event representations has been associated to episodic memory processing and the posterior medial network-PMN, which comprises left angular gyrus-AG, hippocampus, retrosplenial cortex-RSC, parahippocampal formation, posterior cingulate cortex-PCC and anterior medial prefrontal cortex (Ranganath and Ritchey, 2012).

Although PMN is engaged by semantic tasks (Lerner et al., 2011; Silbert et al., 2014), the surge of interest in its role for language is only recent (Hasson et al., 2018). One important question is whether PMN supports construction of semantic representations, besides the core requirement of any situational model to code the spatiotemporal context of the experienced event elements, whose “meaning” might arise from other regions/networks beyond PMN itself. Since activation of episodic information induces activation of semantic representations (and vice versa) (Renoult et al., 2019), it is unclear what modulates PMN’s activity.

To address this question, we undertook new independent analyses on a functional magnetic resonance imaging (fMRI) data set in which participants were presented with written narratives, composed of two consecutive paragraphs: context (1^st^ paragraph) and target (2^nd^ paragraph) (Branzi et al., 2020). We reasoned that construction of an episodic and semantic representation would depend on the number of episodic details and the degree of semantic relatedness between the narrative’s paragraphs, respectively. Separate “episodic richness” and “semantic relatedness” scores were obtained using the Autobiographical Interview (Levine et al., 2002) and Word2vec (Mikolov et al., 2013), respectively. To map the brain regions supporting formation of episodic and semantic representations, these scores were used as parametric modulators of neural responses measured at the onset of the presentation of the 2^nd^ paragraph of each narrative.

Within PMN, RSC may be engaged in the integration of spatiotemporal features presented in a narrative. Instead, left AG and PCC may have a role in integrating semantic information (Ritchey and Cooper, 2020). However, compelling evidence has shown that left AG – particularly the posterior portion (pAG) – and PCC may not be part of the semantic network (Humphreys and Lambon Ralph, 2015). For instance, differently from anterior temporal lobe (ATL) and other parts of the semantic network, left pAG and PCC are deactivated (as compared to rest) in semantic and non-semantic tasks alike (Humphreys and Lambon Ralph, 2017). Instead, the same regions are positively activated by episodic processing tasks (Ritchey et al., 2015; Humphreys et al., 2022b). A possibility is that activity in these regions during previous semantic tasks primarily reflected construction of an episodic (rather than semantic) representation. If correct, neural activity should correlate with the number of episodic details, even after correcting for variance relative to semantic relatedness.

To investigate both key factors that contribute to situation models, we also assessed brain regions primarily involved in processing semantic content. During language, the anterior ventral portion of left AG (avAG) may be engaged in buffering semantic information (Branzi et al., 2021b; Humphreys et al., 2021). If correct, left avAG’s activity should correlate with semantic relatedness, even after correcting for variance relative to episodic richness.

To summarise, we hypothesised that left pAG and PCC would be primarily involved in the processing of episodic information. We expected: given its role in meaning processing (Lambon Ralph et al., 2017), ATL’s activity to show sensitivity to episodic richness; once controlled for semantic relatedness scores, only left pAG and PCC would show strong responses for the construction of episodic representation; RSC’s activity to be modulated by episodic richness only; and semantic relatedness, to modulate ATL’s and possibly left avAG’s activity.

## 2. Materials and Methods

### 2.1. Participants

Participants were the twenty-four volunteers that took part in Branzi et al. (2020). These were 24 adults (19 female), on average aged 23, standard deviation (SD)=3. All participants were right handed (Oldfield, 1971), native English speakers with no history of neurological or psychiatric disorders and normal or corrected-to-normal vision. Because of technical issues during the scanning session, only data from 22 participants (average age=23 years, SD=3; N female=17) were usable for fMRI data analyses. The work described has been carried out in accordance with The Code of Ethics of the World Medical Association (Declaration of Helsinki) for experiments involving humans. Furthermore, all participants gave written informed consent, and the study was approved by the local ethics board.

### 2.2. Stimuli

The stimuli employed in this study were the very same as in Branzi et al. (2020). A total 80 narratives, each one composed by two paragraphs, were used for the experimental study. For each narrative, the same second paragraph (target) was preceded a first paragraph (context) (see Branzi et al., 2020 for the complete list of the stimuli). The present study focussed on the narrative stimuli consisting of textual context and target paragraphs only. There were other types of context and target paragraphs in the original design (see Branzi et al., 2020) (e.g., target paragraphs not preceded by verbal context, number context paragraphs, etc.). However, these were not considered here, because they are irrelevant for the scope of the present study.

### 2.3. Task procedures

The items were presented using an event-related design with the most efficient ordering of events determined using Optseq (http://www.freesurfer.net/optseq). Rest time was intermixed between trials and varied between 2 and 12 seconds (s) (average=3.7, SD=2.8). During this time a red fixation cross was presented to mark the end of each trial (each narrative composed by a context and a target paragraph). Each context paragraph (1^st^ paragraph) was presented at once for 9s followed by the target paragraph (2^nd^ paragraph), which was also presented at once for 6s. A black fixation cross was presented between contexts and targets and its duration varied between 0 and 6s (average=3, SD=1.6). During this time, similarly as during the presentation of the red fixation cross (see above), participants were required to rest.

Participants were asked to read silently both contexts (verbal material and numbers) and targets (only verbal material). Our volunteers were instructed to press a button when arriving to the end of each paragraph (for both contexts and targets). The instruction emphasized speed, but also the need to understand the meaning of verbal contexts and targets since at the end of some of the trials participants would be asked some questions about the content of the narratives. In order to perform this task, it is necessary to integrate the meaning between context and target paragraphs. Hence, following 13% of the trials a comprehension task was presented to ensure that participants were engaged in the task. When this happened, the target item was followed by a statement displayed on the screen for 6s at which participants were required to provide a response (true/false) via button press (see Branzi et al., 2020 for the list of stimuli). A black fixation cross between the target and the comprehension task was presented during a time that varied between 0 and 6s (average=3.5, SD=2.2). Before starting the experimental study, all participants were given written instructions. Then they underwent to a practice session with few trials in order to allow them to familiarise with the task. The stimuli used in the practice session were different from those used in the experimental study.

### 2.4. Task acquisition parameters

Images were acquired using a 3T Philips Achieva scanner using a dual gradient-echo sequence, which is known to have improved signal relative to conventional techniques, especially in areas associated with signal loss (Halai et al., 2014). Thus, 31 axial slices were collected using a TR=2s, TE=12 and 35 milliseconds (ms), flip angle=95°, 80 × 79 matrix, with resolution 3 × 3mm, slice thickness 4mm. For each participant, 1492 volumes were acquired in total, collected in four runs of 746s each.

### 2.5. Data analysis

#### Processing of the narrative stimuli with Autobiographical Interview and Word2vec

Measures of episodic richness were obtained for each narrative stimulus using the Autobiographical Interview method (Levine et al., 2002; Thakral et al., 2017). Thus, we obtained a score which reflected the total number of episodic details (e.g., the contextual who, what, when, where, and why knowledge) contained in each narrative stimulus (**see Figure 1A**).

**Figure 1.**
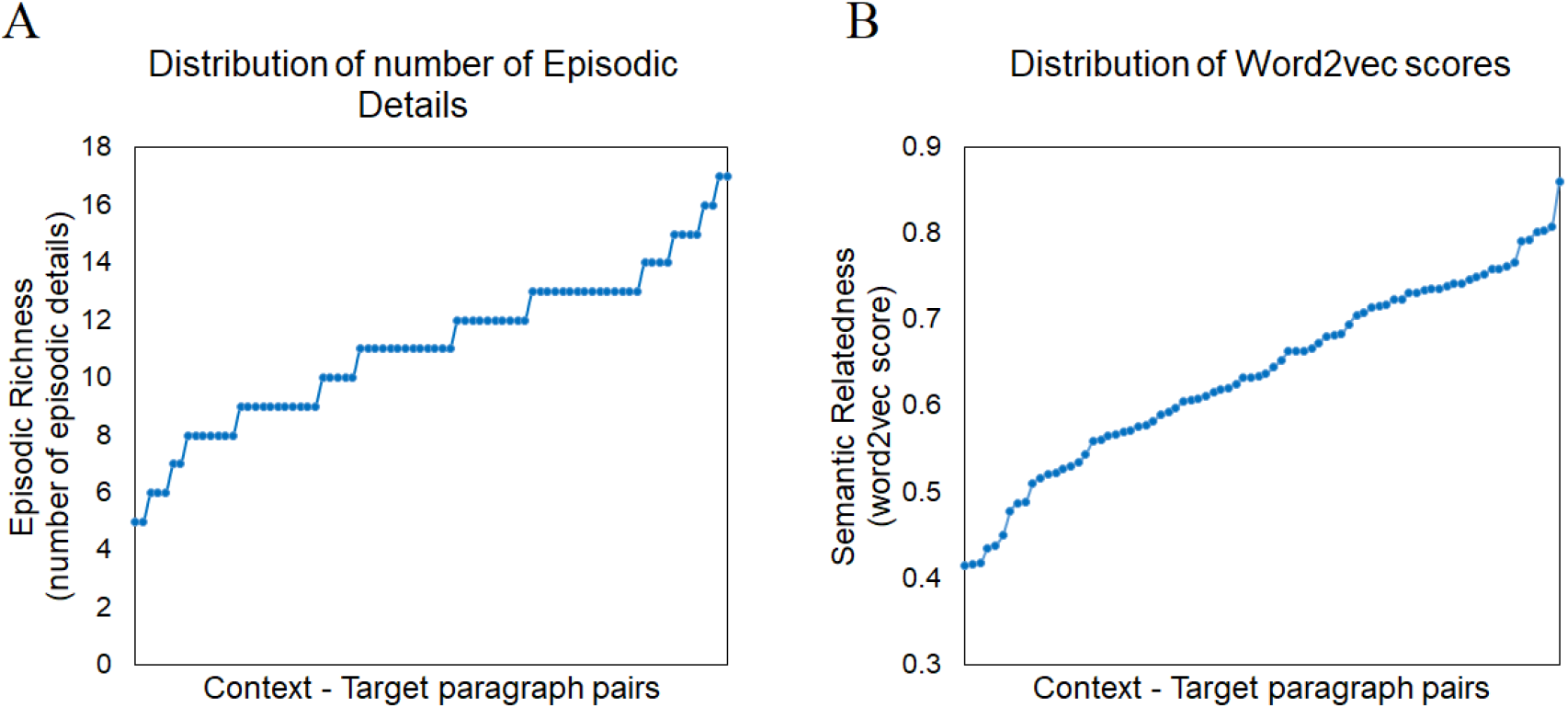
Episodic richness and semantic relatedness scores. Distribution of (A) episodic richness and (B) semantic relatedness scores for all Context-Target paragraph pairs, displayed from lowest to highest.

Measures of semantic relatedness were obtained for each narrative stimulus by subjecting narrative samples to computational analyses based on a pre-trained Google model (https://code.google.com/archive/p/word2vec/). This model provides the user with vector-based representations of the meanings of words, which can be combined linearly to represent the meanings of whole passages of text (e.g., Piai et al., 2016; Hoffman, 2019). For each narrative we obtained two vectors representing the semantic content of context and target paragraphs. Then, for each narrative the context and target vectors were compared using a cosine similarity metric (the cosine gives a value for similarity between zero and one), allowing us to obtain for each narrative a score reflecting the between-paragraphs semantic relatedness (or coherence) (**see Figure 1B**).

A high score indicates that the semantic representation of the context-paragraph is closely related to that of the target-paragraph, whereas a low coherence value indicates that the semantic content of the context paragraph is less related to the topic being presented in the target paragraph. Analyses were implemented in Python (the code is publicly available at http://www.mrc-cbu.cam.ac.uk/publications/opendata/). Preprocessing of the data included the removal of stop-words (high frequency words which do not add much meaning to a sentence, e.g., articles, pronouns) using the genism library, as well as the exclusion of some words which could not be found in the model (“Rubik” and “Elsa”).

#### fMRI data analyses

##### Preprocessing

The dual-echo images were averaged. Data were analysed using SPM12. After motion-correction images were co-registered to the participant’s T1 image. Spatial normalisation into MNI space was computed using DARTEL (Ashburner, 2007), and the functional images were resampled to a 3 × 3 × 3mm voxel size and smoothed with an 8mm FWHM Gaussian kernel.

#### Whole brain analyses

The data were filtered using a high-pass filter with a cut-off of 128s and then analysed using a General Linear Model (GLM). We ran a GLM in which, at the individual subject level, each condition was modelled with a separate regressor (target paragraphs) with time derivatives added, and events were convolved with the canonical hemodynamic response function, starting from the onset of the 2^nd^ paragraph (target paragraph). The context paragraphs (1^st^ paragraphs), the target paragraphs preceded by nonverbal (numerical) contexts, and the comprehension task were also modelled as regressors. However, these conditions were not further analysed because they are not relevant for the scope of the present study. Each condition was modelled as a single event with a duration corresponding to 6s (target conditions and comprehension task trials) or 9s (context conditions). Motion parameters were entered into the model as covariates of no interest. A voxel-height threshold of *p* < .001 was adopted for all whole-brain analyses, with correction for multiple comparisons performed at the cluster level (family wise error (FWE) corrected at *p* < .05).

#### Activation varying with episodic richness and semantic relatedness

To determine the brain regions modulated by episodic richness (score reflecting the number of episodic details) and semantic relatedness (Word2vec score), we conducted two separate whole-brain analyses. In these analyses, both episodic and semantic scores were used as parametric modulators of the neural responses measured at the onset of the presentation of the 2^nd^ paragraph. In respect to the modelling of these neural responses, we were expecting neural responses to change linearly as a function of episodic and semantic content. Nevertheless, neural responses reflecting an accumulation of semantic and episodic information might not necessarily increase in a linear fashion (Hoffman, 2019). Therefore, we also examined non-linear (quadratic and cubic function) models.

For both episodic and semantic parametric modulators, the same polynomial expansions were applied to investigate both linear and non-linear relationships (whole-brain regression analyses) between the blood-oxygen-level-dependent imaging (BOLD) response and these parameters. Importantly, since scores for episodic richness and semantic relatedness were positively correlated (*r* = 0.430, *p* < .001), and since we examined linear and non-linear responses which are required to be orthogonalised in the model, we adopted one recommended solution for our analyses. That is, we employed two models where the parametric modulators of interest were orthogonalised (e.g., Mumford et al., 2015), so that all other modulators coming after the first one (e.g., episodic richness) were orthogonalised with respect to the first (e.g., semantic relatedness) in a stepwise function. Thus, in one model, to assess the neural correlates of episodic richness, we specified semantic relatedness and episodic richness as the first and second parametric modulator, respectively. Instead, in the other model, to assess the neural correlates of semantic relatedness, episodic richness was the first parametric modulator of the target condition, whilst semantic relatedness was the second parametric modulator. We also examined the neural correlates of episodic richness and semantic relatedness without correction for semantic and episodic variables, respectively. This is important to determine the extent to which the observed functional dissociations may depend on the orthogonalisation procedure.

## 3. Results

### Activation varying with episodic richness and semantic relatedness

#### The neural correlates of episodic richness

The results revealed both linear and nonlinear relationships between neural activity measured during narrative processing and the parameter reflecting episodic content. Aligning with our hypothesis, neural activity measured in AG (avAG and pAG), the ATL and PCC/Precuneus increased linearly with the number of episodic details presented in the narratives (see **Figure 2A** and **Table 1**). Interestingly and divergent to the ATL and PCC/Precuneus, AG’s activity (bilaterally) was significantly modulated by episodic content, even after controlling for semantic relatedness. The results also revealed a nonlinear significant relationship (inverse cubic function) between neural activity measured in the RSC (bilateral) and episodic richness (see **Figure 2B, 2C** and **Table 1)**.

**Figure 2.**
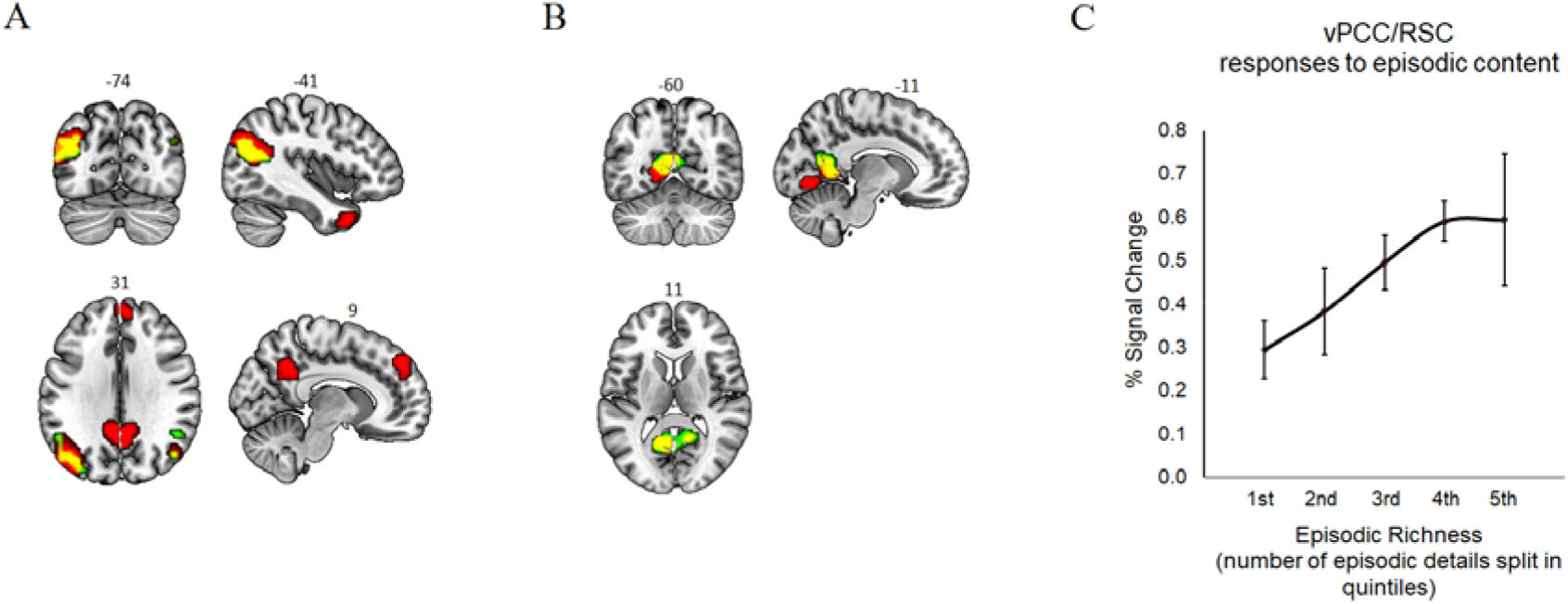
Neural correlates of episodic richness. Whole-brain results reflect the (A) linear and (B) nonlinear (inverse cubic function) relationship between the number of episodic details in the narratives and the neural activity measured during narrative processing (the statistical maps were voxel-wise thresholded at *p* < .001, and whole-brain cluster extent FWE-corrected at *p* < .05). Red = results from the model where the variance relative to semantic relatedness was *not* partialled out; Green = results from the model where the variance relative to semantic relatedness was partialled out; Yellow = overlap between the two models. (C) The plot shows the non-linear effect of episodic richness on neural activity measured in ventral (v) PCC/RSC (% signal change). The mask corresponding to the vPCC/RSC was identified from the whole-brain results (vPCC/RSC cluster). Episodic scores were divided into five sets based on their values (where 1st quintile = from 5 to 8 and 5th quintile = from 14 to 17). This type of averaging is used for illustration purposes only. Bars indicate standard error of the mean.

**Table 1.**
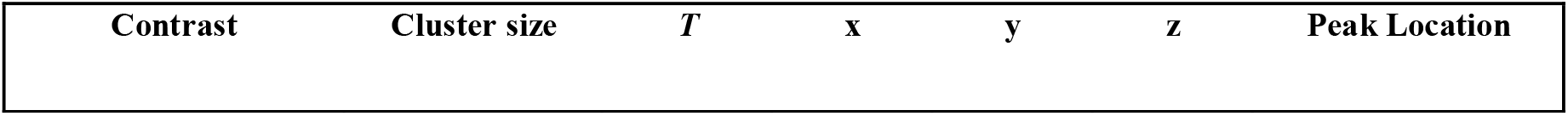
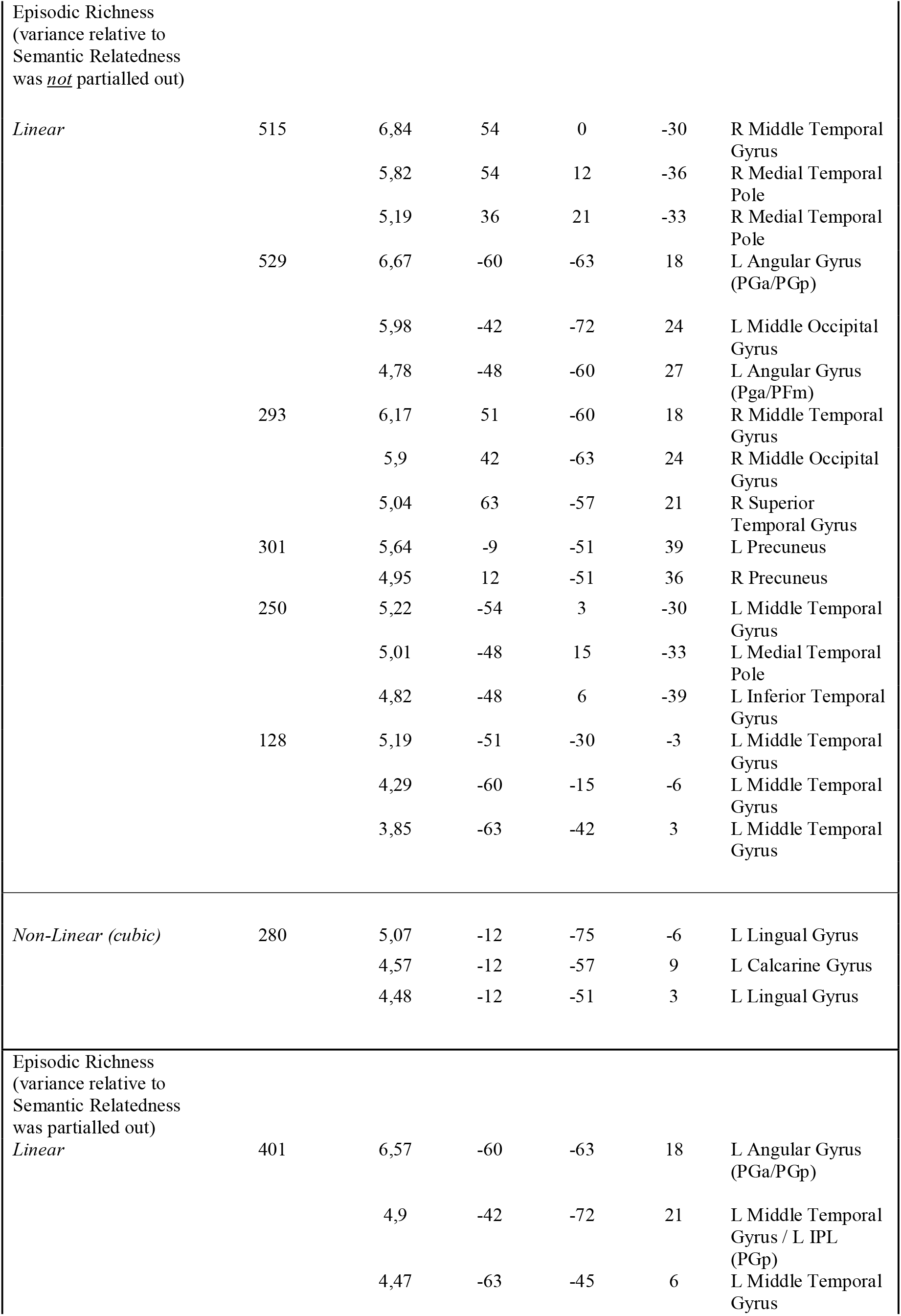
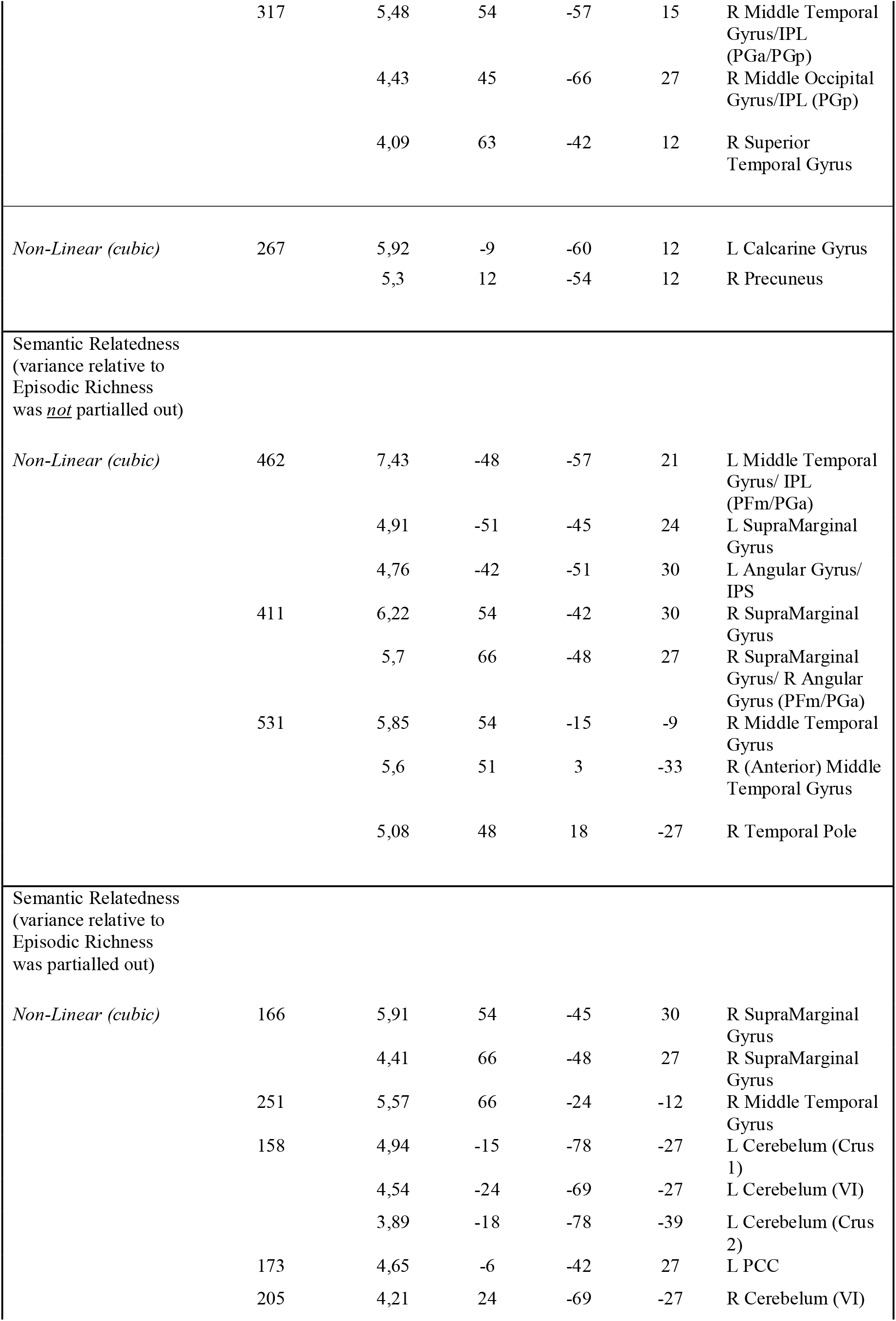
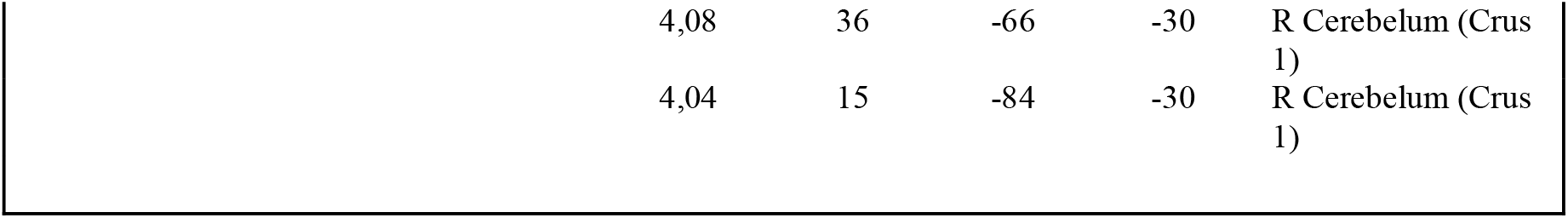
The locations of the activation peaks (MNI coordinates) from the whole-brain analyses where statistical maps were voxel-wise thresholded at *p* < .001, and whole-brain cluster extent FWE-corrected at *p* < .05. *Abbreviations*: L = Left; R= Right; PCC = Posterior Cingulate Cortex; IPL = Inferior Parietal Lobe; IPS = intra-parietal sulcus.

#### The neural correlates of semantic relatedness

The results revealed a significant nonlinear relationship (inverse cubic function) between neural activity measured during narrative processing and semantic relatedness scores. These modulations were observed in different regions within the PMN (see **Figure 3A** and **Table 1**). Unlike the left avAG, the activity in right ATL/middle temporal gyrus (MTG), left dorsal PCC (dPCC), and right inferior parietal lobe, including both supramarginal gyrus (SMG) and avAG, was significantly modulated by semantic relatedness, even after correcting for the variance relative to episodic richness. A plot of the neural responses in right ATL/MTG revealed an overall positive relationship between activity in this region and semantic relatedness scores (see **Figure 3B**).

**Figure 3.**
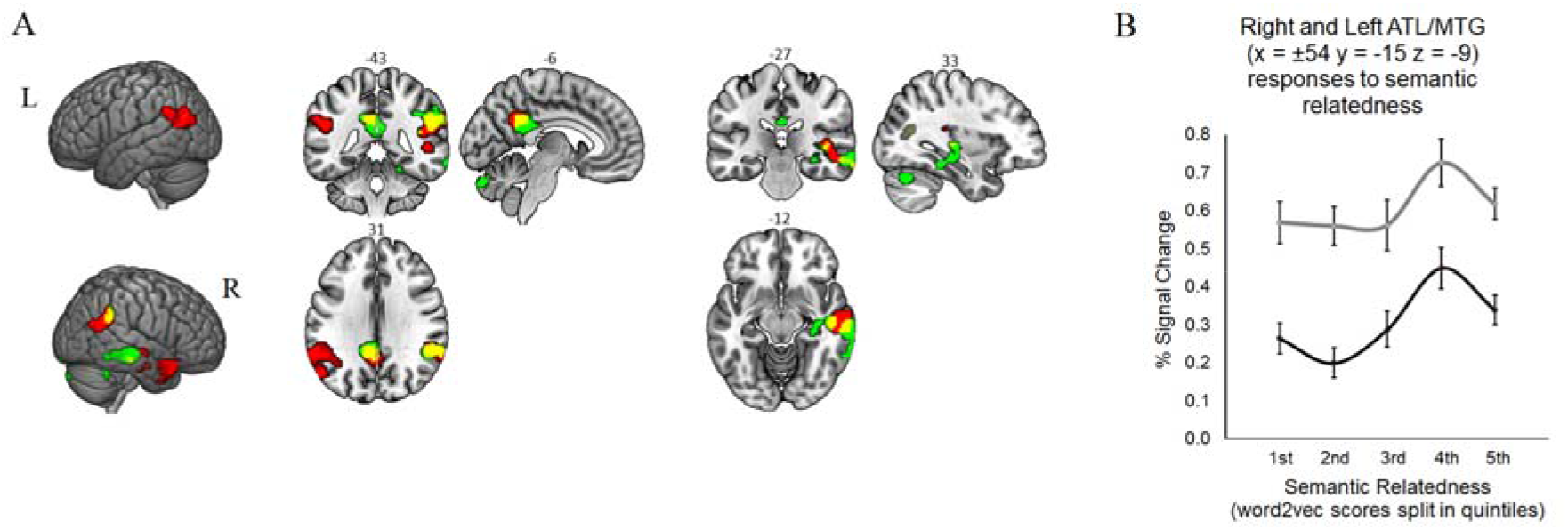
Neural correlates of semantic relatedness. (A) Whole-brain results reflect the nonlinear (inverse cubic function) relationship between semantic relatedness between the paragraphs of the narrative and the neural activity measured during narrative processing (the statistical maps were voxel-wise thresholded at *p* < .001, and whole-brain cluster extent FWE-corrected at *p* < .05). Red = results from the model where the variance relative to episodic richness was *not* partialled out; Green = results from the model where the variance relative to episodic richness was partialled out; Yellow = overlap between the two models. (B) The black line in the plot reflects the non-linear effect of semantic relatedness on neural activity measured in right ATL/MTG (% signal change). For this region of interest (ROI) analysis, a 6mm radius sphere centered at x = 54 y = −15 z = −9 was employed. This sphere was overlapping with the right ATL/MTG cluster modulated by semantic relatedness (whole-brain results). For visual comparison, we also report neural responses (grey line) in the controlateral ROI in the left hemisphere (6mm radius sphere centered at x = −54 y = −15 z = −9). Semantic relatedness scores were divided into five sets based on their values (where 1st quintile = least coherent and 5th quintile = most coherent). This type of averaging is used for illustration purposes only. Bars indicate standard error of the mean.

Importantly, some brain areas (e.g., hippocampus in **Figure 3A**) were modulated by semantic relatedness only in the model where the two parametric modulators were orthogonalised. We refrain to interpret these results since they may be due to effects of no interest of the first parametric modulator on the second one. For instance, it could be that the first parametric modulator affects the second by reducing the residual error or by suppressing other variables in the model.

Considering the results for each parameter together (see **Figure 4** and **Table 1**), we found that the left AG (both and anterior ventral and posterior portions) and ventral (v) PCC/RSC support the construction of the episodic content of the narratives. In contrast, the right ATL, right avAG/SMG, left dPCC, support the formation of the semantic content of the narratives. Importantly, the episodic *versus* semantic dissociation observed in the ventral and dorsal PCC respectively, is unlikely to be due to the orthogonalization of the parameters. In fact, as it can be observed in Figures 2 and 3, the dissociation is present also in the models without orthogonalisation of the parameters.

**Figure 4.**
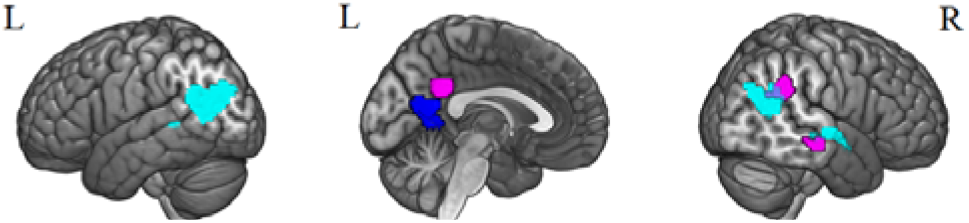
Summary of results: The unique neural correlates of semantic relatedness and episodic richness. This figure shows the neural overlap between models with and without orthogonalization (Yellow blobs in Figures 2 and 3). In *cyan* whole-brain results reflecting the positive linear relationship between episodic richness and neural activity. In *blue* whole-brain results reflecting the nonlinear (inverse cubic function) relationship between episodic richness and neural activity. In *purple* whole-brain results reflecting the nonlinear (inverse cubic function) relationship between semantic relatedness and neural activity. All the statistical maps were voxel-wise thresholded at *p* < .001, and whole-brain cluster extent FWE-corrected at *p* < .05.

## 4. Discussion

The construction of complex mental representations reflecting current events is a key feature of many daily activities, including language comprehension. Whilst there is a consensus that these representations involve the combination of both semantic and episodic information, it is still unclear whether the same neural regions, particularly those within the PMN, support the formation of semantic or episodic content, or both (Renoult et al., 2019). In the present fMRI study, we addressed this question using a narrative reading task. We extracted from the narrative stimuli both semantic and episodic measures, and then we correlated them with neural activity. This analysis allowed us to establish which PMN regions support the construction of semantic representations independently from their episodic content, and vice versa.

### Construction of episodic representations

We hypothesised that left AG – particularly the posterior portion – and PCC would be primarily involved in the construction of episodic representations, and thus show neural responses proportional to the number of episodic details (e.g., Thakral et al., 2017, 2020). We also hypothesised that episodic richness would modulate activity measured in RSC (for a review see Ritchey and Cooper, 2020). Indeed, we found that left AG, including posterior and anterior ventral portions, was sensitive to the episodic content of the narratives, in accord with previous studies (e.g., Baldassano et al., 2017; van der Linden et al., 2017; Humphreys et al., 2022a, 2022b). The positive activation profiles observed in left AG, together with the observation that activity within this region is sensitive to the amount of episodic information, provide strong support for the interpretation that left AG supports online maintenance/buffering of episodic information (Vilberg and Rugg, 2008; Humphreys and Lambon Ralph, 2015; Humphreys et al., 2022b) and accords with a recent transcranial magnetic stimulation study in which we showed that the disruption of this AG-online buffering process during language processing impairs formation of memory associations between the episodic details presented in a narrative (Branzi et al., 2021b).

We also found that activity in ventral (v) PCC/RSC was modulated by episodic richness. These results align with a substantial literature suggesting a key role for this region in episodic memory (Valenstein et al., 1987; Rudge and Warrington, 1991; Gainotti et al., 1998; Aggleton and Pearce, 2001; Vincent et al., 2006; Epstein et al., 2007). For instance, patients with focal lesions to vPCC/RSC show amnestic syndromes (Valenstein et al., 1987; Heilman et al., 1990; Rudge and Warrington, 1991; Takayama et al., 1991; Katai et al., 1992; Gainotti et al., 1998; McDonald et al., 2001). Furthermore, RSC and surrounding PCC are affected since the initial stages of Alzheimer’s disease, which is characterised by episodic encoding deficits (Nestor et al., 2003). This collective, convergent evidence points to a prominent role of vPCC/RSC in episodic construction processes. Indeed, this region has strong reciprocal connections with the hippocampal complex via the cingulum bundle (Morris et al., 1999; Kobayashi and Amaral, 2003, 2007), and evidence suggests that RSC would act as a mediating hub between more dorsal parts of PMN and episodic encoding systems (parahippocampal formation/hippocampus) (Kaboodvand et al., 2018). Thus, one interesting possibility is that, in the present study, episodic details are continuously buffered by left AG (online maintenance of episodic representations reflected by sustained neural responses) and then, these representations are slowly integrated in the hippocampus (perhaps in event-chunks), via the RSC ‘mediating hub’.

### Construction of semantic representations

We were expecting semantic relatedness to modulate neural activity within the PMN, and particularly the activity of left avAG (Price et al., 2015; van der Linden et al., 2017; Jackson, 2020). Beyond the PMN, we also expected the activity of ATL to be modulated by the semantic content of the narratives, given ATL’s prominent role in semantic cognition (Lambon Ralph et al., 2017).

We found that left avAG and ATL were both modulated by semantic relatedness. However, when the variance relative to episodic richness was partialled out, the semantic modulation effect in left avAG disappeared. This result suggests that this region may not have a key role in the construction of semantic coherent representations per se (e.g., Humphreys et al., 2021). Activity in right parietal cortex (SMG/avAG), instead, was significantly modulated by the semantic content, along with that of other regions, including left dPCC and right ATL/MTG.

Our results linked the activity of the dPCC to the formation of coherent semantic representations. This result accords with evidence suggesting that PCC responds to the coherence of event structure (Hasson et al., 2008; Lerner et al., 2011; Simony et al., 2016) and that PCC representations generalise across auditory and visual modalities (Baldassano et al., 2017). From those studies, however, it was not clear the extent to which this generalisation involved semantic rather than episodic content. Our study shows that the anterior dorsal portion of the PCC, but not the vPCC/RSC, may be involved in the construction of complex semantic gist-like representations. This result, which is unlikely to reflect a task difficulty effect (reading times and Word2vec scores were uncorrelated: *r* = 0.165, *p* = .145), is consistent with evidence showing that whilst vPCC/RSC is preferentially involved in representing spatiocontextual features (e.g., Epstein, 2008; Bicanski and Burgess, 2018), dPCC may be tuned towards nonspatial, conceptual information (Horner et al., 2015; Baldassano et al., 2017; Silson et al., 2019). If so, it is unclear why PCC does not appear to be consistently activated by semantic tasks (Noonan et al., 2013; Humphreys et al., 2015; Krieger-Redwood et al., 2015; Canini et al., 2016; Jackson, 2020; Branzi et al., 2021a; Jackson et al., 2021). One possibility is that its role becomes relevant when the semantic task requires formation of multi-item complex gist-like representations, i.e., during time-extended and context-dependent semantic tasks. Since these types of tasks are not typically included in these large-scale semantic meta-analyses (Noonan et al., 2013; Jackson, 2020), it is not surprising that this brain region has not been regularly identified as a potential key area of the semantic network. If dPCC supports the building of an evolving semantic representation, one important question to be addressed in the future is whether dPCC and ATL/MTG represent the same type of semantic information during language processing.

A second somewhat unexpected finding refers to the lateralization of the neural effects observed for semantic relatedness. For instance, we found a clear modulation of semantic relatedness in the right but not in the left ATL/MTG. Similarly, our results show that the right avAG/SMG especially tracks semantic coherence. One possibility is that the left hemisphere is already strongly involved regardless of the semantic content, and that the right hemisphere becomes primarily involved only when information presented in the second paragraph is highly coherent with the previous context. This explanation is in accord with our results showing a stronger engagement of left *versus* right ATL/MTL, irrespective of the semantic content (see **Figure 3B**). Another possibility is that left and right ATL/MTG are supporting semantic processes operating at different time scales. Whilst it is well-established that human language functions are mostly lateralized to the left hemisphere of the brain, it has been also suggested that the right hemisphere, as compared to the left, may be especially involved for processing global (sentence or paragraph), rather than local (single words) aspects of linguistic contents (St George et al., 1999; Hickok and Poeppel, 2007; Poeppel, 2003). Since the semantic scores used in this study reflected semantic relatedness across paragraphs, it may not be surprising to find the right, but not the left, ATL/MTG to respond to semantic relatedness.

## Conclusions

Our results confirm that parts of the PMN, traditionally associated to episodic memory processing, play a key role also in language comprehension and semantic processing. These findings may allow a better understanding of linguistic deficits in patients with episodic memory impairments and represent a first step towards novel investigations on the interaction between episodic and semantic memory systems during language processing. Future studies will have to address whether the episodic *versus* semantic neural dissociations here observed generalise also to task involving nonverbal modalities (nonverbal visual and auditory modalities), as well as *where* and *how* episodic and semantic information is integrated during language comprehension.

## References

Aggleton JP, Pearce JM (2001) Neural systems underlying episodic memory: insights from animal research. Philos Trans R Soc Lond B Biol Sci 356:1467–1482.

Ashburner J (2007) A fast diffeomorphic image registration algorithm. Neuroimage 38:95–113.

Bailey HR, Kurby CA, Sargent JQ, Zacks JM (2017) Attentional focus affects how events are segmented and updated in narrative reading. Mem Cognition 45:940–955.

Baldassano C, Chen J, Zadbood A, Pillow JW, Hasson U, Norman KA (2017) Discovering Event Structure in Continuous Narrative Perception and Memory. Neuron 95:709–721 e705.

Bicanski A, Burgess N (2018) A neural-level model of spatial memory and imagery. Elife 7.

Binder JR, Desai RH (2011) The neurobiology of semantic memory. Trends Cogn Sci 15:527–536.

Branzi FM, Martin CD, Paz-Alonso PM (2021a) Task-Relevant Representations and Cognitive Control Demands Modulate Functional Connectivity from Ventral Occipito-Temporal Cortex During Object Recognition Tasks. Cereb Cortex.

Branzi FM, Humphreys GF, Hoffman P, Lambon Ralph MA (2020) Revealing the neural networks that extract conceptual gestalts from continuously evolving or changing semantic contexts. Neuroimage:116802.

Branzi FM, Pobric G, Jung J, Lambon Ralph MA (2021b) The Left Angular Gyrus Is Causally Involved in Context-dependent Integration and Associative Encoding during Narrative Reading. J Cogn Neurosci:1–14.

Canini M, Della Rosa PA, Catricala E, Strijkers K, Branzi FM, Costa A, Abutalebi J (2016) Semantic interference and its control: A functional neuroimaging and connectivity study. Hum Brain Mapp 37:4179–4196.

Epstein RA (2008) Parahippocampal and retrosplenial contributions to human spatial navigation. Trends in Cognitive Sciences 12:388–396.

Epstein RA, Parker WE, Feiler AM (2007) Where am I now? Distinct roles for parahippocampal and retrosplenial cortices in place recognition. J Neurosci 27:6141–6149.

Gainotti G, Almonti S, Di Betta AM, Silveri MC (1998) Retrograde amnesia in a patient with retrosplenial tumor. Neurocase 4:519–526.

Halai AD, Welbourne SR, Embleton K, Parkes LM (2014) A comparison of dual gradient-echo and spin-echo fMRI of the inferior temporal lobe. Hum Brain Mapp 35:4118–4128.

Hasson U, Egidi G, Marelli M, Willems RM (2018) Grounding the neurobiology of language in first principles: The necessity of non-language-centric explanations for language comprehension. Cognition 180:135–157.

Hasson U, Yang E, Vallines I, Heeger DJ, Rubin N (2008) A hierarchy of temporal receptive windows in human cortex. J Neurosci 28:2539–2550.

Heilman KM, Bowers D, Watson RT, Day A, Valenstein E, Hammond E, Duara R (1990) Frontal Hypermetabolism and Thalamic Hypometabolism in a Patient with Abnormal Orienting and Retrosplenial Amnesia. Neuropsychologia 28:161–169.

Hickok G, Poeppel D (2007) The cortical organization of speech processing. Nat Rev Neurosci 8:393–402.

Hoffman P (2019) Reductions in prefrontal activation predict off-topic utterances during speech production. Nat Commun 10:515.

Horner AJ, Bisby JA, Bush D, Lin WJ, Burgess N (2015) Evidence for holistic episodic recollection via hippocampal pattern completion. Nat Commun 6.

Humphreys GF, Lambon Ralph MA (2015) Fusion and Fission of Cognitive Functions in the Human Parietal Cortex. Cereb Cortex 25:3547–3560.

Humphreys GF, Lambon Ralph MA (2017) Mapping Domain-Selective and Counterpointed Domain-General Higher Cognitive Functions in the Lateral Parietal Cortex: Evidence from fMRI Comparisons of Difficulty-Varying Semantic Versus Visuo-Spatial Tasks, and Functional Connectivity Analyses. Cereb Cortex 27:4199–4212.

Humphreys GF, Lambon Ralph MA, Simons JS (2021) A Unifying Account of Angular Gyrus Contributions to Episodic and Semantic Cognition. Trends Neurosci 44:452–463.

Humphreys GF, Halai AD, Branzi FM, Lambon Ralph MA (2022). The angular gyrus is engaged by autobiographical recall not object-semantics, or event-semantics: Evidence from contrastive propositional speech production. bioRxiv 2022.04.04.487000; doi: https://doi.org/10.1101/2022.04.04.487000.

Humphreys GF, Jung J, Lambon Ralph MA (2022b) The convergence and divergence of episodic and semantic functions across lateral parietal cortex. Cereb Cortex.

Humphreys GF, Hoffman P, Visser M, Binney RJ, Lambon Ralph MA (2015) Establishing task- and modality-dependent dissociations between the semantic and default mode networks. Proc Natl Acad Sci U S A 112:7857–7862.

Jackson RL (2020) The neural correlates of semantic control revisited. Neuroimage 224:117444.

Jackson RL, Rogers TT, Lambon Ralph MA (2021) Reverse-engineering the cortical architecture for controlled semantic cognition. Nat Hum Behav 5:774–786.

Kaboodvand N, Backman L, Nyberg L, Salami A (2018) The retrosplenial cortex: A memory gateway between the cortical default mode network and the medial temporal lobe. Human Brain Mapping 39:2020–2034.

Katai S, Maruyama T, Hashimoto T, Yanagisawa N (1992) [A case of cerebral infarction presenting as retrosplenial amnesia]. Rinsho Shinkeigaku 32:1281–1287.

Kobayashi Y, Amaral DG (2003) Macaque monkey retrosplenial cortex: II. Cortical afferents. J Comp Neurol 466:48–79.

Kobayashi Y, Amaral DG (2007) Macaque monkey retrosplenial cortex: III. Cortical efferents. J Comp Neurol 502:810–833.

Krieger-Redwood K, Teige C, Davey J, Hymers M, Jefferies E (2015) Conceptual control across modalities: graded specialisation for pictures and words in inferior frontal and posterior temporal cortex. Neuropsychologia 76:92–107.

Lerner Y, Honey CJ, Silbert LJ, Hasson U (2011) Topographic mapping of a hierarchy of temporal receptive windows using a narrated story. J Neurosci 31:2906–2915.

Levine B, Svoboda E, Hay JF, Winocur G, Moscovitch M (2002) Aging and autobiographical memory: Dissociating episodic from semantic retrieval. Psychol Aging 17:677–689.

McDonald CR, Crosson B, Valenstein E, Bowers D (2001) Verbal encoding deficits in a patient with a left retrosplenial lesion. Neurocase 7:407–417.

Mikolov T, Chen K, Corrado G, Dean J (2013) Efficient estimation of word representations in vector space. arXiv preprint arXiv:1301.3781.

Morris R, Petrides M, Pandya DN (1999) Architecture and connections of retrosplenial area 30 in the rhesus monkey (Macaca mulatta). Eur J Neurosci 11:2506–2518.

Mumford JA, Poline JB, Poldrack RA (2015) Orthogonalization of Regressors in fMRI Models. Plos One 10.

Nestor PJ, Fryer TD, Ikeda M, Hodges JR (2003) Retrosplenial cortex (BA 29/30) hypometabolism in mild cognitive impairment (prodromal Alzheimer’s disease). Eur J Neurosci 18:2663–2667.

Noonan KA, Jefferies E, Visser M, Lambon Ralph MA (2013) Going beyond Inferior Prefrontal Involvement in Semantic Control: Evidence for the Additional Contribution of Dorsal Angular Gyrus and Posterior Middle Temporal Cortex. J Cognitive Neurosci 25:1824–1850.

Oldfield RC (1971) The assessment and analysis of handedness: the Edinburgh inventory. Neuropsychologia 9:97–113.

Piai V, Anderson KL, Lin JJ, Dewar C, Parvizi J, Dronkers NF, Knight RT (2016) Direct brain recordings reveal hippocampal rhythm underpinnings of language processing. Proc Natl Acad Sci U S A 113:11366–11371.

Poeppel D (2003) The analysis of speech in different temporal integration windows: cerebral lateralization as ‘asymmetric sampling in time’. Speech Commun 41:245–255.

Price AR, Bonner MF, Peelle JE, Grossman M (2015) Converging evidence for the neuroanatomic basis of combinatorial semantics in the angular gyrus. J Neurosci 35:3276–3284.

Lambon Ralph MA, Jefferies E, Patterson K, Rogers TT (2017) The neural and computational bases of semantic cognition. Nat Rev Neurosci 18:42–55.

Ranganath C, Ritchey M (2012) Two cortical systems for memory-guided behaviour. Nature Reviews Neuroscience 13:713–726.

Renoult L, Irish M, Moscovitch M, Rugg MD (2019) From Knowing to Remembering: The Semantic-Episodic Distinction. Trends in Cognitive Sciences 23:1041–1057.

Ritchey M, Cooper RA (2020) Deconstructing the Posterior Medial Episodic Network. Trends in Cognitive Sciences 24:451–465.

Ritchey M, Montchal ME, Yonelinas AP, Ranganath C (2015) Delay-dependent contributions of medial temporal lobe regions to episodic memory retrieval. Elife 4.

Rudge P, Warrington EK (1991) Selective Impairment of Memory and Visual-Perception in Splenial Tumors. Brain 114:349–360.

Silbert LJ, Honey CJ, Simony E, Poeppel D, Hasson U (2014) Coupled neural systems underlie the production and comprehension of naturalistic narrative speech. Proc Natl Acad Sci U S A 111:E4687–4696.

Silson EH, Steel A, Kidder A, Gilmore AW, Baker CI (2019) Distinct subdivisions of human medial parietal cortex support recollection of people and places. Elife 8.

Simony E, Honey CJ, Chen J, Lositsky O, Yeshurun Y, Wiesel A, Hasson U (2016) Dynamic reconfiguration of the default mode network during narrative comprehension. Nat Commun 7:12141.

St George M, Kutas M, Martinez A, Sereno MI (1999) Semantic integration in reading: engagement of the right hemisphere during discourse processing. Brain 122 (Pt 7):1317–1325.

Takayama Y, Kamo H, Ohkawa Y, Akiguchi I, Kimura J (1991) [A case of retrosplenial amnesia]. Rinsho Shinkeigaku 31:331–333.

Thakral PP, Madore KP, Schacter DL (2017) A Role for the Left Angular Gyrus in Episodic Simulation and Memory. Journal of Neuroscience 37:8142–8149.

Thakral PP, Madore KP, Schacter DL (2020) The core episodic simulation network dissociates as a function of subjective experience and objective content. Neuropsychologia 136.

Valenstein E, Bowers D, Verfaellie M, Heilman KM, Day A, Watson RT (1987) Retrosplenial Amnesia. Brain 110:1631–1646.

van der Linden M, Berkers R, Morris RGM, Fernandez G (2017) Angular Gyrus Involvement at Encoding and Retrieval Is Associated with Durable But Less Specific Memories. J Neurosci 37:9474–9485.

Vilberg KL, Rugg MD (2008) Memory retrieval and the parietal cortex: a review of evidence from a dual-process perspective. Neuropsychologia 46:1787–1799.

Vincent JL, Snyder AZ, Fox MD, Shannon BJ, Andrews JR, Raichle ME, Buckner RL (2006) Coherent spontaneous activity identifies a hippocampal-parietal memory network. J Neurophysiol 96:3517–3531.

Zwaan RA, Langston MC, Graesser AC (1995) The construction of situation models in narrative comprehension: An event-indexing model. Psychological Sci 6: 292–297.

Zwaan RA, Radvansky GA (1998) Situation models in language comprehension and memory. Psychol Bull 123:162–185.

